# Building an Open Representation for Biological Protocols

**DOI:** 10.1101/2022.07.05.498808

**Authors:** Bryan Bartley, Jacob Beal, Miles Rogers, Daniel Bryce, Robert P. Goldman, Benjamin Keller, Peter Lee, Vanessa Biggers, Joshua Nowak, Mark Weston

**Affiliations:** Raytheon BBN Technologies, USA; SIFT, LLC, USA; University of Washington, USA; Ginkgo Bioworks, USA; Strateos, Inc., USA; Netrias, Inc., USA

**Keywords:** protocol, biology, representation, UML, RDF, SBOL

## Abstract

Laboratory protocols are critical to biological research and development, yet difficult to communicate and reproduce across projects, investigators, and organizations. While many attempts have been made to address this challenge, there is currently no available protocol representation that is unambiguous enough for precise interpretation and automation, yet simultaneously abstract enough to enable reuse and adaptation. The Protocol Activity Markup Language (PAML) is a free and open protocol representation aiming to address this gap, building on a foundation of UML, Autoprotocol, and SBOL RDF. PAML provides a representation both for protocols and for records of their execution and the resulting data, as well as a framework for exporting from PAML for execution by either humans or laboratory automation. PAML is currently implemented in the form of an RDF knowledge representation, specification document, and Python library, can be exported for execution as either a manual “paper protocol” or Autoprotocol, and is being further developed as an open community effort.

## 1 INTRODUCTION

Laboratory protocols are critical to biological research and development. However, protocols are often difficult to communicate or reproduce, given the differences in context, skills, instruments, and other resources between different projects, investigators, and organizations. One of the necessary preconditions for effectively addressing these challenges is for there to be at least one commonly used data representation for describing laboratory protocols that is unambiguous enough for precise interpretation and automation, yet simultaneously abstract enough to support reuse and adaptation.

While there has been much prior work on representations for protocols, prior approaches have generally been limited either by their dependence on natural language or in the expressiveness of their representation. Many protocol representations focus on simplifying the capture and distribution of descriptions in natural language, such as protocols.io [27] and the many commercial electronic laboratory notebook products. A similar approach is used for recording protocol execution information with community-defined minimum information standards such as MIAME [7], MIFlowCyt [14], and STRENDA [28]. Much of the key information in such approaches is encoded using natural language, which is easier to solicit from experimentalists, but cannot be readily interpreted by machines. As a consequence, protocols and protocol execution records captured with such representations cannot be automatically validated and are often ambiguous, incorrect, or lacking key information. For the same reasons, such protocols cannot generally be executed by laboratory automation systems.

Other protocol representations have focused specifically on automation-assisted execution. In many cases, these are highly specific solutions tied to specific hardware, often proprietary and tied to particular vendors. Some representations have been made applicable to a broader set of automation systems, however, such as Autoprotocol [17] and Antha [25], or instead use laboratory technicians as their automation, as in the case of Aquarium [12]. All of these representations, however, have been generally “low level” in their description of protocols, focusing on very specific details of each operation. This specificity, which on the one hand enables automated execution, on the other hand poses barriers to adoption and to generalization and reuse, since this level of detail often obscures understanding and is too tied to the specifics of a particular laboratory to readily transfer into different environments. Such representations are typically also difficult to translate into more “human-friendly” forms.

Finally, there are a number of workflow languages that solve similar problems in business logic or information processing, such as UML [19], the Common Workflow Language [2], Taverna [30], and Cromwell [8], not to mention biology-specific workflow systems such as Toil [29] and Galaxy [11]. The execution models of such systems are in some cases are general enough to be applied to the description and execution of laboratory protocols, but to the best of our knowledge such an application has not previously been implemented. Further, their very generality can make it more difficult for a domain expert to see how to apply them to protocols, given the large gap between abstract task execution concepts and the specifics of particular tasks that must be performed in a laboratory.

Here, we present a unified approach to protocol representation that addresses all of these disparate needs and bridges prior approaches through the development of the Protocol Activity Markup Language (PAML), a free and open protocol representation building on a foundation of UML [19], Autoprotocol [17], and SBOL3 RDF [3, 16]. In Section 2, we elaborate on the design goals for PAML, its three foundations, and the approach taken for implementation. Section 3 outlines the key elements of PAML’s representation for protocols, libraries of primitive actions, and execution records. Section 4 then discusses the current prototype implementation of PAML and the execution environments it supports, and Section 5 discusses ongoing plans for development.

## 2 ARCHITECTURE

We begin by elaborating a minimal set of design requirements necessary for any broadly applicable representation of biological protocols. These requirements led us to identify a set of core representational ingredients sufficient to address these requirements.

### 2.1 Design Requirements

Information about laboratory protocols is used for a wide range of purposes in research and development, at many different stages of experiment design, execution, data analysis, interpretation, and communication and sharing with other groups. As such, to be effective as a broadly shared community standard, we argue that any protocol representation will need to be able to support at least the following goals:

- **Execution by either humans or machines:** When available, laboratory automation can greatly improve the productivity of researchers, so protocols should be specified in sufficient detail to enable them to be mapped for machine execution. Many laboratories, however, do not have automation available. Moreover, even when some degree of automation is available, it is common for protocols to incorporate both automated and manual stages, so protocols also need to be able to be presented in a succinct and human-friendly form.
- **Maintaining execution records and associated metadata markup:** When a protocol is actually executed, it is important to be able to record the specific time of execution, the laboratory and personnel that executed the protocol, equipment used, etc. A protocol representation thus needs to include support for creating persistent data structures that record a specific protocol execution and linking such execution records back to protocol specifications. It is also important to be able to support automation in metadata tagging of the data collected in the course of protocol execution. For example, given a protocol that collects flow cytometry data on samples of different strains under varying growth conditions, the protocol specification should support automatic association/markup linking each FCS file with information about the strain, growth conditions, time of data collection, calibration, etc. of the sample from which the FCS file was produced.
- **Mapping protocols from one laboratory environment to another:** Protocol replication and reuse requires the ability to map a protocol from one laboratory to another, despite their differences in the specific equipment, inventory, and information systems. A protocol representation cannot guarantee that a protocol can be transferred, particularly one that is poorly understood or delicate in execution. Rather, a specification should allow a protocol to say, to the best of the authors’ knowledge, how to predict if a mapping will product a correct execution and how to check if an execution should be considered correct, i.e., what a protocol specification truly requires, which aspects can safely be varied, which must be honored, and what are the anticipated tolerances for inputs and outputs.
- **Recording modifications of protocols and the relationship between different versions:** Protocols are likely to be the subject of ongoing improvement and maintenance. For example, a protocol may be modified in order to enable the protocol to be simpler to execute or more reliable, to be executed at a lab with different equipment from the lab at which the protocol originated, to enable the protocol to be scaled up or scaled down, etc. A likely use pattern is to have a protocol initially be “too strict” (too specific) to be instantiated in a new lab that wishes to run the protocol, and thus need modification. Rather than creating a new protocol, ideally, the original should be generalized to allow it to run both where it could before and also in the new lab. This improved version could then be contributed back and released as an updated version of the existing protocol. Alternatively, if for some reason generalization is impossible or impractical, users should be able to create a variant and record the source of that variant.
- **Verification and validation of protocol completeness and coherence:** Authoring a protocol requires substantial care and effort, and the usefulness of the protocol can be compromised if its specification is ill-formed, erroneous, or incomplete (e.g., the classic “inadequate methods section” issue in scientific publications). Supporting protocol authors in achieving correctness is thus an important goal for a protocol representation, and while the implementation will depend on specific tooling, the representation specification must provide guidance as to what it means for a protocol to be complete, consistent, etc. This is especially important for automatically-executed protocols since the control system cannot be counted on to repair flaws in protocols on the fly, and in the worst case, an incorrect specification could even cause damage to equipment or endanger lab personnel.
- **Planning, scheduling, and allocation of laboratory resources:** Laboratory resources are valuable, and some organizations will want to be able to optimize their use. To do so, a protocol representation should support (at least) extraction of resource requirements and estimated durations from activities in the protocol. Note that the specifics of resource requirements and duration estimates will likely be a function of both the protocol and the available equipment in the laboratory in which it is to be executed. Which resources are limited, and must be considered in a planner or scheduler, versus those that can be effectively treated as unlimited, will also vary by laboratory, as will management styles and applicable policies.

### 2.2 Foundations: UML, Autoprotocol, and SBOL RDF

In developing the PAML protocol representation, we adopted a principle of building upon existing standards wherever possible, in order to increase compatibility and interoperability, take advantage of existing tooling, and make the implementation as lightweight as possible. We introduce these foundations here at a conceptual level, while the specifics of their usage are provided in Section 3.

#### 2.2.1 UML Behavior Models

As the core of a protocol is a workflow of activities to be carried out, we began by identifying an established standard for workflow modeling that could provide both a well-defined and general formal semantics, yet also be sufficiently abstract as to allow succinct expression and adaptation. We found such a model in Unified Modeling Language (UML) behavior representations (specifically the current version 2.5.1 [19]). UML behaviors provide a general, domain-independent workflow model. This model encodes a formal execution model based on token-passing, which can support serial, parallel, non-deterministic, and distributed execution. Furthermore, as UML use often focuses on diagram-based communication, it also provides a set of flow control and abstraction constructs for succinct and human-friendly communication about complex workflows. At the same time, the formal execution model provides an unambiguous semantics for verification, validation, and other forms of machine reasoning.

#### 2.2.2 Autoprotocol Laboratory Primitives

As UML is domain-independent, it does not provide any guidance on what primitive behaviors are suitable for expressing laboratory-independent behaviors. For this, we turn to Autoprotocol [17]. Autoprotocol describes biological protocols in terms of a sequence of instructions, and while this linear workflow is not expressive enough for our requirements, the instructions themselves are primitives that can be readily mapped from one laboratory environment to another. These instructions, such as `liquid_handle`, incubate, provision, and spin have been specifically designed and refined by the authors of Autoprotocol as a basis set for expressing the activities of common biological protocols in a manner readily transported between different pieces of laboratory automation. The Autoprotocol instructions thus provide a reasonable starting point for our library of laboratory-independent primitive behaviors.

Some activities expressed using these primitives are at a lower level than desirable for a human experimenter, however, such as specifying pipette mixing as a sequence of repetitive liquid handling operations. In cases such as these, we will choose to not use Autoprotocol instructions as defined, but replace them with more complex or abstract alternatives that instead capture an Autoprotocol use pattern.

#### 2.2.3 SBOL 3 Materials, Records, and RDF

While UML models processes, it does not actually provide a representation to capture executions and associate traces with data. For this, we turn to the Synthetic Biology Open Language (SBOL), version 3 [3, 16], which uses Semantic Web practices and resources, such as *Uniform Resource Identifiers* (URIs) and ontologies, to unambiguously identify and define biological system elements and to provide serialization formats for encoding this information in electronic data files.

While the early versions of SBOL focused only on genetic designs, it has since been expanded to represent and link information throughout the design-build-test-learn workflow [16]. SBOL provides succinct representations for all of the materials that would be used by a typical biological protocol—strains, reagents, media, experimental sample designs, etc.—along with the ability to track and distinguish between specific physical aliquots and replicates. On the input side of a protocol, SBOL’s combinatorial design specifications [23] offer the ability to compactly specify combinations of experimental conditions. SBOL also incorporates the Ontology of Units of Measure (OM) [22] for specifying and recording measurements, as well as the W3C Provenance Ontology (PROV-O) [18] for linking specifications, samples, and data via traces of activity records.

This last is precisely complementary to our selection of UML behaviors for representing workflows, as PROV-O leaves the actual definition of activities to users. Recall that a key objective of PAML is to aid in tracing the connections between data sets and the protocols that produced them—the issue of data set provenance. The provenance ontology (PROV-O) is a well-accepted tool for encoding this kind of information, so we propose to use it to anchor the data-producing relationship between protocols, executions of protocols, and the resulting data sets. PROV-O also specifies several annotation properties that we adopt to mark up protocol executions. Thus, we can use PROV-O as the basis for capturing execution traces, with the activities in the trace defined using the UML data model, built from laboratory primitives based on Autoprotocol, and the inputs, outputs, and data relations encoded using SBOL 3.

SBOL’s semantic web basis has also been used to allow representational extensions with custom classes without requiring changes to the underlying specification, unlike UML or Autoprotocol. For this reason, we select SBOL RDF as the underlying data model for PAML and convert the relevant portions of UML and Autoprotocol into SBOL RDF extensions. This approach also allows PAML to be extended with additional custom information for particular uses and deployments.

Critically, SBOL RDF provides a partial closure reasoning model (see Section 5.5 of [3]) that allows much stronger and more “object-oriented” reasoning than plain RDF or OWL, while still allowing documents to reference external material. This allows for an intuitive chunking and linking of information, for example, being able to store and reason about a complete record of a protocol execution that has links to the protocol without being required to store a copy of the protocol in the same document.

Finally, SBOL RDF also offers a natural approach to management of protocol modifications and versioning, since SBOL RDF can be serialized into the sorted N-triples RDF format. This format is a stable serialization that can be readily differenced and inspected with standard, text-based version control tooling. Thus, implementing PAML using SBOL RDF allows protocols to be maintained by distributed communities of contributors using standard software development version control such as git, as well as the larger ecosystem of associated tooling for project management and community-driven development.

## 3 PAML DATA MODEL

Following the architecture presented in Section 2, the PAML data model is formulated as an extension of SBOL 3, implemented by encoding an ontology for the UML behavior model and its supporting classes, plus additional classes for linking this information into PROV-O records, libraries of primitive UML activities based on Autoprotocol, and additional classes for tracking laboratory samples and data. In this section, we present all of the major concepts and classes of PAML; additional supporting classes and the complete current specification can be found in the PAML draft specification at http://bioprotocols.org.

Following the pattern of SBOL, PAML’s data model is defined in terms of **classes**, instantiated as **objects**. Classes support **inheritance** based on subclass and superclass relationships. Under the SBOL RDF partial closure model, all objects inherit from the `sbol:Identified` type, which supports generic identity and annotation information. Objects that describe a complete subsystem for reasoning additionally inherit from the `sbol:TopLevel` type, and are always bundled with their “child” `sbol:Identified` objects, allowing strong closed-world reasoning over any collection of `sbol:TopLevel` objects. Examples of `sbol:TopLevel` classes in PAML are `paml:Protocol` and `paml:BehaviorExecution`, while examples of child classes are `uml:ActivityNode` and `uml:Parameter`, which are only useful in the context of the `paml:Protocol` they are used to describe.

Classes contain data in the form of **properties**, which may have primitive XML types (string, integer, long, float, boolean, URI, etc.) or may refer to another object via its URI; the former are **data properties**, and the latter **object properties**. Diagrams for the data model use UML conventions, in which classes are shown as rectangles labelled at the top with their class name (or containing nothing but the class name for classes defined elsewhere), and an arrow with a hollow triangle at its head represents the inheritance relationship between subclass (tail) and superclass (head). Data properties are shown in text within the rectangle, object properties shown via an arrow with a diamond at the end, and class inheritance by arrows with empty heads. For object properties, filled diamonds indicate **association properties**, in which an object “owns” a child object: this is important because it means the child object is bundled with its parent under SBOL RDF’s partial closure model and always moves with its parent in an RDF document to enable strong (closed-world) reasoning about its contents. Empty diamonds, on the other hand, indicate references to objects defined elsewhere, which may or may not be available to reason about without retrieving additional documents. Finally, the number of values a property can have is indicated by upper and lower points on its **cardinality**: [1] (shorthand for 1… 1]) indicates a required property, [0 … 1] indicates an optional property, [0 · · · *] (the empty constraint) indicates a property that can have any number of values, and [1… *] a property that must have at least one value.

For clarity, since there are several ontologies involved in the implementation of the PAML data model and at least pair of important terms from different ontologies with the same shortened name (`uml:Activity` vs. `prov:Activity`), we include ontology prefixes for all terms in the text. Likewise, in the figures presenting the data we color model classes by ontology and include their ontology prefix; properties are not, however, as they always belong to the ontology of their defining class (e.g., `uml:ownedParameter` is a property for `uml:Behavior`).

To illustrate the data model, we will use the iGEM LUDOX protocol for calibration of plate reader optical density (OD) measurements [24], first introduced in the 2016 iGEM interlaboratory study [6] and refined thereafter [5]. This is an extremely simple protocol consisting of three steps: water is added to four wells in a 96-well plate, LUDOX silica suspension is added to another four wells, and then all eight samples are measured at some specified absorbance (600nm in its original usage), in order to obtain a baseline measurement of OD, validate machine behavior, and allow path-length correction.

### 3.1 Protocols

In UML 2.5.1 [19]), the basic building block of process modeling is a `uml:Behavior`, which is an abstract specification for how the state of a system changes over time. A `uml:Activity` is a type of `uml:Behavior` that defines a process in terms of a network of steps linked together by flows of information and control. For PAML, we import a strict subset of UML 2.5.1 into SBOL RDF, omitting those classes and properties that we do not need. Note that this does not require any modification of the SBOL standard, as these are implemented as extension classes outside of the restricted SBOL namespace and conform with the SBOL specification guidance on extension via custom classes.

#### 3.1.1 Laboratory Primitives are UML Behaviors

Figure 1 shows the class structure for the adaptation of `uml:Behavior` and `uml:Activity` into SBOL RDF. A UML `uml:Behavior` provides the definition an interface for a process. The `uml:ownedParameter` property links to an ordered list of `uml:Parameter` objects (each marked internally with a direction), which describe the order and type of the input arguments that can be given when the `uml:Behavior` is invoked and of the output values that will be returned when the `uml:Behavior` completes its execution. Additional optional `uml:precondition` and `uml:postcondition` properties provide `uml:Constraint` objects that specify requirements on `uml:Parameter` values before and after execution, respectively.

**Fig. 1.**
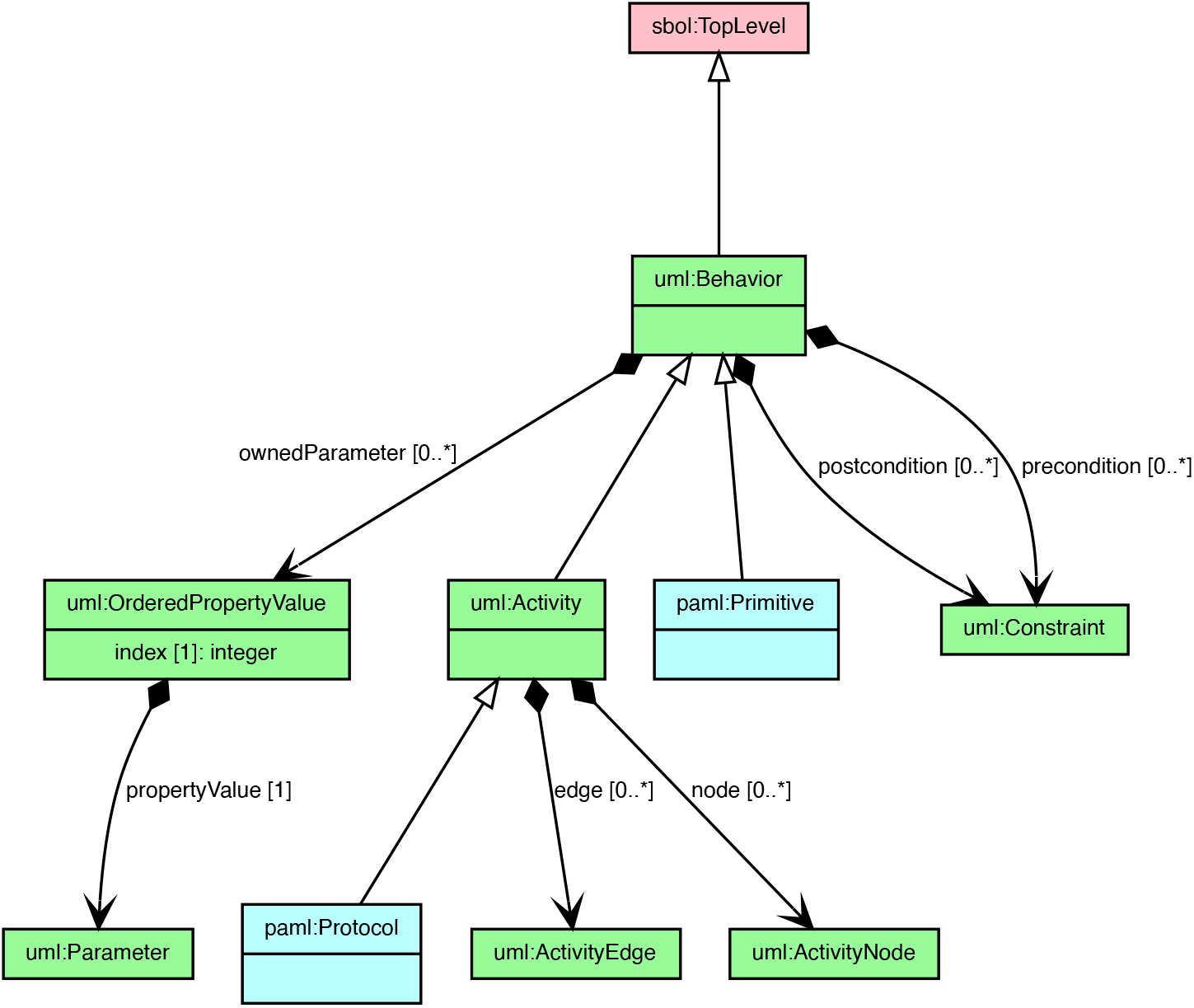
The `uml:Activity` and `uml:Behavior` classes. A `paml:Protocol` is defined as a `uml:Activity`, while a `paml:Primitive` laboratory action is defined as a `uml:Behavior`.

In PAML, the simplest protocol building block is a `paml:Primitive`, defined as a subclass of UML `uml:Behavior`. Primitives are basic laboratory operations such as pipetting, measuring absorbance in a plate reader, or centrifuging. For example, the iGEM LUDOX calibration protocol uses a `paml:Primitive` named `paml:Provision` to dispense water and LUDOX into a plate and another named `paml:MeasureAbsorbance` to measure the OD values of these samples (both of these are adapted from Autoprotocol as descibed below in Section 3.1.3).

Primitives do not provide any information about how to carry out the corresponding real world action (such as pipetting). Instead, they serve as the handoff point between PAML and an execution environment that knows how to actually carry out such primitives in a laboratory, as described further in Section 4.3. However, the *constraints* on these primitives can be used specify what is expected to happen (how the system changes) when the corresponding actions are performed.

In order to build useful protocols in PAML, we also need libraries of primitives that are simple enough to be readily reused, yet abstract enough to be easily transferred from laboratory to laboratory. As noted above, Autoprotocol [17] already provides a good beginning set of primitive operations, which have already been validated as both reusable and transferrable between different pieces of laboratory equipment. In adapting Autoprotocol, however, PAML adds two key extensions to make protocols more adaptable and reusable: classes for describing collections of samples and classes for organizing primitive operations into libraries.

#### 3.1.2 Sample Collections

In Autoprotocol it is only possible to address one location at a time, i.e., a single container or a single compartment within a container, such as a well on a plate. This means that any operation on multiple locations must name every location as a fixed value in the definition of the protocol, which in turn means that protocols are often both extremely large (e.g., individually operating on every well of a 96-well plate) and rigid, since locations cannot be supplied as a parameter value or determined dynamically at runtime.

In common practice, however, protocols are often naturally described in terms of operations on physical or logical collections of samples, such as “Wells A1 to D2 in a standard 96-well plate,” “A 6 by 3 group of 10ml tubes: three replicates each for six conditions,” or “All wells showing green fluorescence >500 MEFL.” Being able to represent such descriptions directly allows representations to be more compact, more intelligible to humans for both authoring and execution, and also more reusable, since they can be communicated as parameters or determined dynamically. PAML thus includes a `paml:SampleCollection` class, as shown in Figure 2(a), that represents organized collections of samples, either as an n-dimensional `paml:SampleArray` (e.g., the wells of a 96-well plate, a sequence of 10 flasks) or as a `paml:SampleMask` that selects a subset of such an array using a mask of Boolean values.

**Fig. 2.**
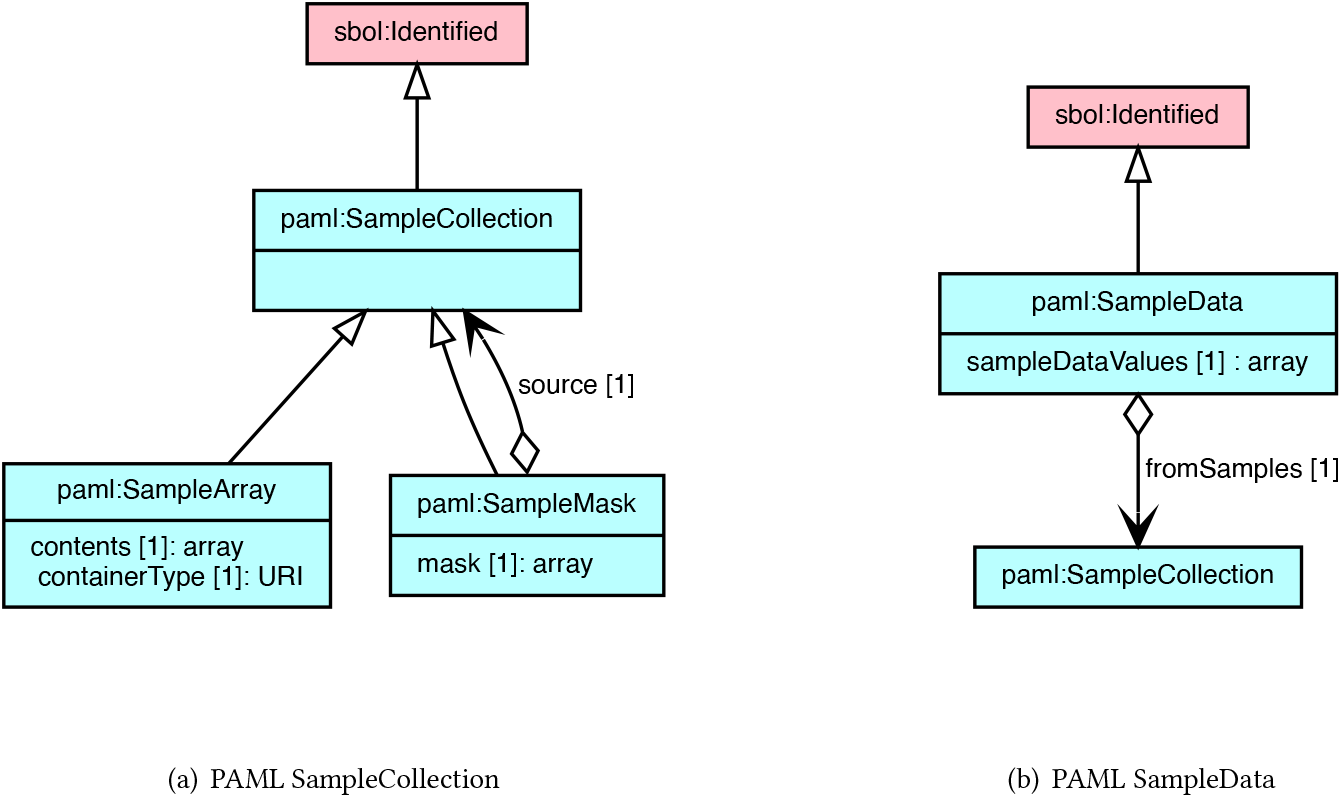
PAML represents collections of samples as either N-dimensional arrays or using Boolean masks to select subsets from such an array (a). PAML represents sample data by associating an array of data values with a collection of samples (b).

In particular, A `paml:SampleArray` specifies an n-dimensional rectangular array of samples, all stored in the same type of container. For example, in the iGEM LUDOX calibration protocol, a `paml:SampleArray` is generated by the `paml:EmptyContainer` primitive to allocate a new 96-well plate for use in the protocol. A `paml:SampleArray` might also be used to describe a set of 10 cell cultures growing in 96-well plate wells, or a set of 6 streaked agar plates, or a single 500 mL flask filled with media. For any non-empty location, the contents are in turn described by an SBOL 3 `sbol:Implementation` object that represents a physical sample and which, in turn, can link to an SBOL 3 `sbol:Component` object that describes the mixture of materials in that location. Note that this is a logical array, and does not necessarily indicate the actual layout of the samples in space, which may be determined by context in the execution environment. For example, a 2×4 array of samples in 96-well plate wells might end up being laid out as a 2×4 array in wells A1 to B4 or as a 2×4 array in wells G5 to H8 or as an 8×1 column in wells A1 to H1, or even as eight wells scattered arbitrarily around the plate according to an anti-bias quality control schema. This also allows for higher-dimensional arrays where each dimension represents an experimental factor. For example, an experiment testing four factors with 3, 3, 4, and 5 values per factor, for a total of 180 combinations, could be represented as a 4-dimensional `sbol:sample` array of 96-well plate wells, and then end up laid out over two plates.

As an alternative to a sample array, a `paml:SampleMask` describes a subset of samples in some `paml:SampleCollection` (array or mask) by an array of Boolean values, where true values indicate that a sample is included and false values indicate that it is excluded. To allow masks to be readily composed and interchanged, their dimension is kept identical to that of the source `paml:SampleCollection`. In this way, references to the physical laboratory elements on which a `paml:SampleArray` is laid out can be easily maintained and combined even across different subsetting operations. For example, in the iGEM LUDOX calibration protocol, a `paml:SampleMask` is used to select wells A1 to D1 on the plate to fill with water and another `paml:SampleMask` used to select wells A2 to D2 to fill with LUDOX, then a third `paml:SampleMask` covers both with the range A1 to D2 for measuring absorbance.

Finally, PAML defines another class, `paml:SampleData`, as shown in Figure 2(b), in order to capture the relationship between physical samples and information about those samples. The `paml:SampleData` class simply associates a `paml:SampleCollection` with an array that has values defined for all of the included samples of the collection, thereby representing measurements, such as an array of plate reader absorbance measurements. The joining of samples and data with the `paml:SampleData` class also allows information to be fed back into the computation of `paml:SampleMask` objects at runtime, such as the example above of “all wells showing green fluorescence >500 MEFL.”

In sum, with the addition of the `paml:SampleCollection` and `paml:SampleData` classes, PAML allows primitives to be defined in terms of operations on collections of samples, rather than on specific locations, e.g., dispensing media into a collection of wells or measuring the absorbance in those wells. While there are some cases, such as our simple example of the iGEM LUDOX calibration protocol, where specific locations are appropriate, a great many protocols are intended to be able to run over a set of samples or conditions that are provided as input. PAML primitives are thus in general defined for the more general case of collections, with references to specific locations being just one of the ways in which such a collection may be defined.

#### 3.1.3 Primitive Libraries

PAML also adds libraries for organizing collections of primitives. In Autoprotocol there is a fixed set of primitives defined by the specification, which limits extensibility. For PAML, we take an approach like that used in most modern programming languages, in which the specified language is kept as small as possible, while many of the capabilities of the language are provided by collections of functions grouped into libraries, each library associated with a well-defined cluster of functionality.

Figure 3 shows how the current implementation of PAML organizes operations taken from Autoprotocol into four libraries of `paml:Primitive` objects. Three of the libraries are classes of laboratory activities that operate on `paml:SampleCollection` objects. The `paml:plate_handling` library contains operations performed on containers (e.g., plates, flasks), such as sealing and incubation, which are mostly direct mapping of equivalent Autoprotocol activities. The `paml:liquid_handling` library contains operations moving liquids within or between containers, such as pipetting from one location to another or using a pipette to mix fluids in a location. This includes a division of the omnibus Autoprotocol `liquid_handle` operation into several different patterns of usage, which we have chosen to do in order to make higher-level abstractions that are both more readily human-interpretable and also more readily accessible for machine reasoning and verification. The third library, `paml:spectrophotometry`, does the same for plate reader measurements. Finally, we have implemented one library that contains mostly new operations (i.e., with no Autoprotocol equivalents) for creating and subsetting `paml:SampleCollection` objects. A few Autoprotocol operations are not included in the current implementation, but are expected to be adapted to add into existing or new libraries as implementation continues.

**Fig. 3.**
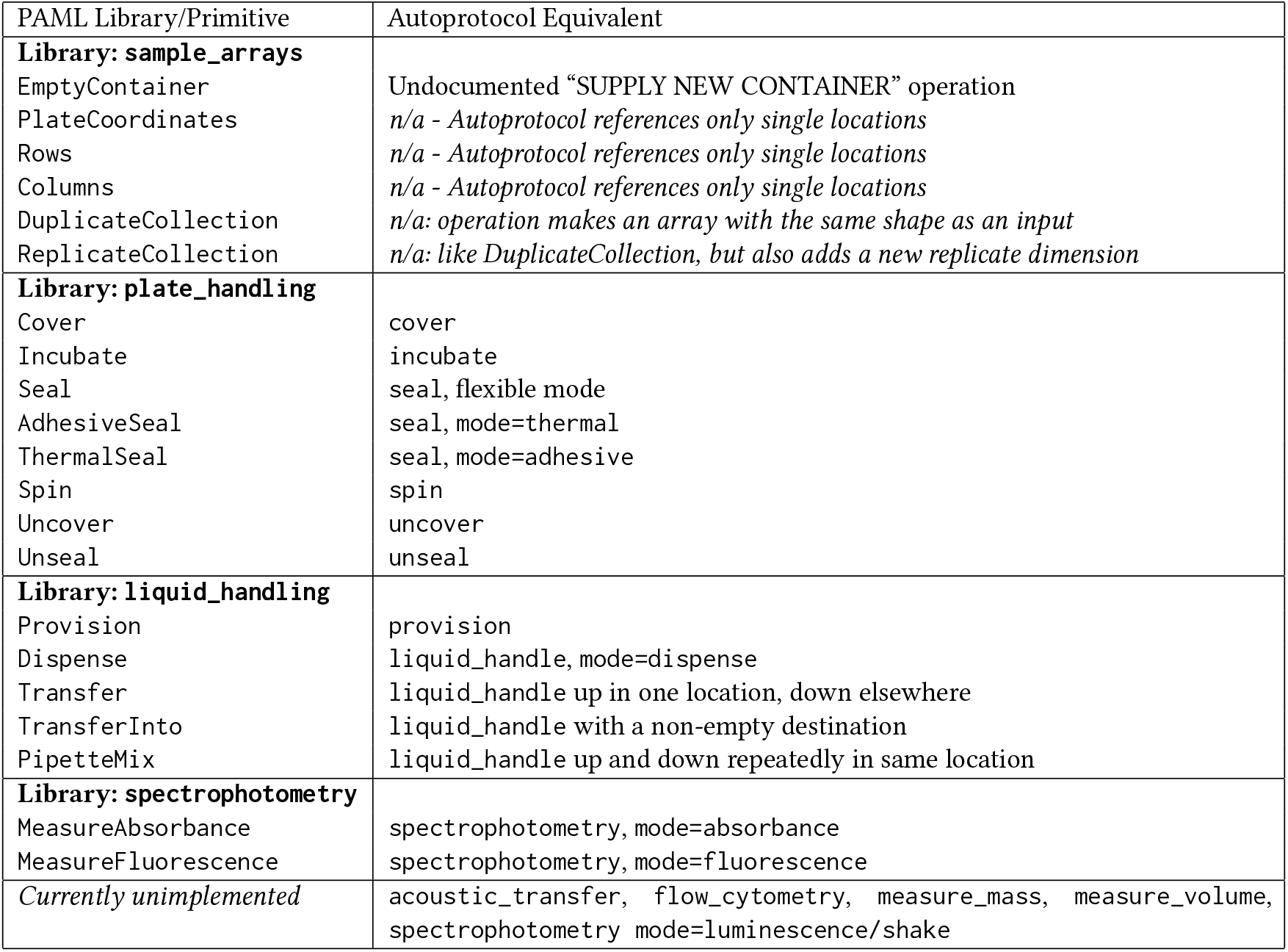
PAML’s primitive laboratory operations are organized into libraries based on required equipment types. The initial collection of “built-in” primitives are based on Autoprotocol, currently implementing most functionality from that language, plus primitives for defining and manipulating collections of samples.

Organizing primitives into libraries and aligning libraries with equipment provides a basis for comparing protocol requirements and laboratory capabilities. A protocol’s capability requirements may be coarsely determined by the set of libraries that it uses. Any given protocol execution environment can then be defined in terms of which libraries are supported. Moreover, different libraries may be supported with different means of execution. For example, a laboratory with a liquid handling robot and a plate reader may support automated execution of `paml:Primitive` operations from the `paml:liquid_handling` library, while those in `paml:plate_handling` and `paml:spectrophotometry` are carried out by a human operator.

#### 3.1.4 Protocols are UML Activities

We now move from individual laboratory operations to complete protocols. In UML, a `uml:Activity` is used to define composite behaviors in terms of a network of steps. Its `uml:node` property stores a collection of `uml:ActivityNode` objects (Figure 4(a)). In the current implementation of PAML, these `uml:ActivityNodes` can only be of three sub-types:

1. A `uml:CallBehaviorAction` object defines an actual step of the protocol, in which a specific `paml:Primitive` or sub-`paml:Protocol` is carried out (note that this class is several layers deep in a hierarchy of other sibling UML `uml:ExecutableNode` classes not currently used in PAML).
2. A `uml:ObjectNode` defines a point where data enters and exits the `uml:Activity` (by means of a `uml:Activity`ParameterNode) or one of its `uml:CallBehaviorAction` nodes (by means of a `uml:Pin`).
3. A `uml:ControlNode` is used to define where the steps of the `uml:Acitivity` start (`uml:InitialNode`), stop (`uml:FinalNode`), branch (splitting at a `uml:DecisionNode` and rejoining at a `uml:MergeNode`), or run in parallel (splitting at a `uml:ForkNode` and rejoining at a `uml:JoinNode`).

**Fig. 4.**
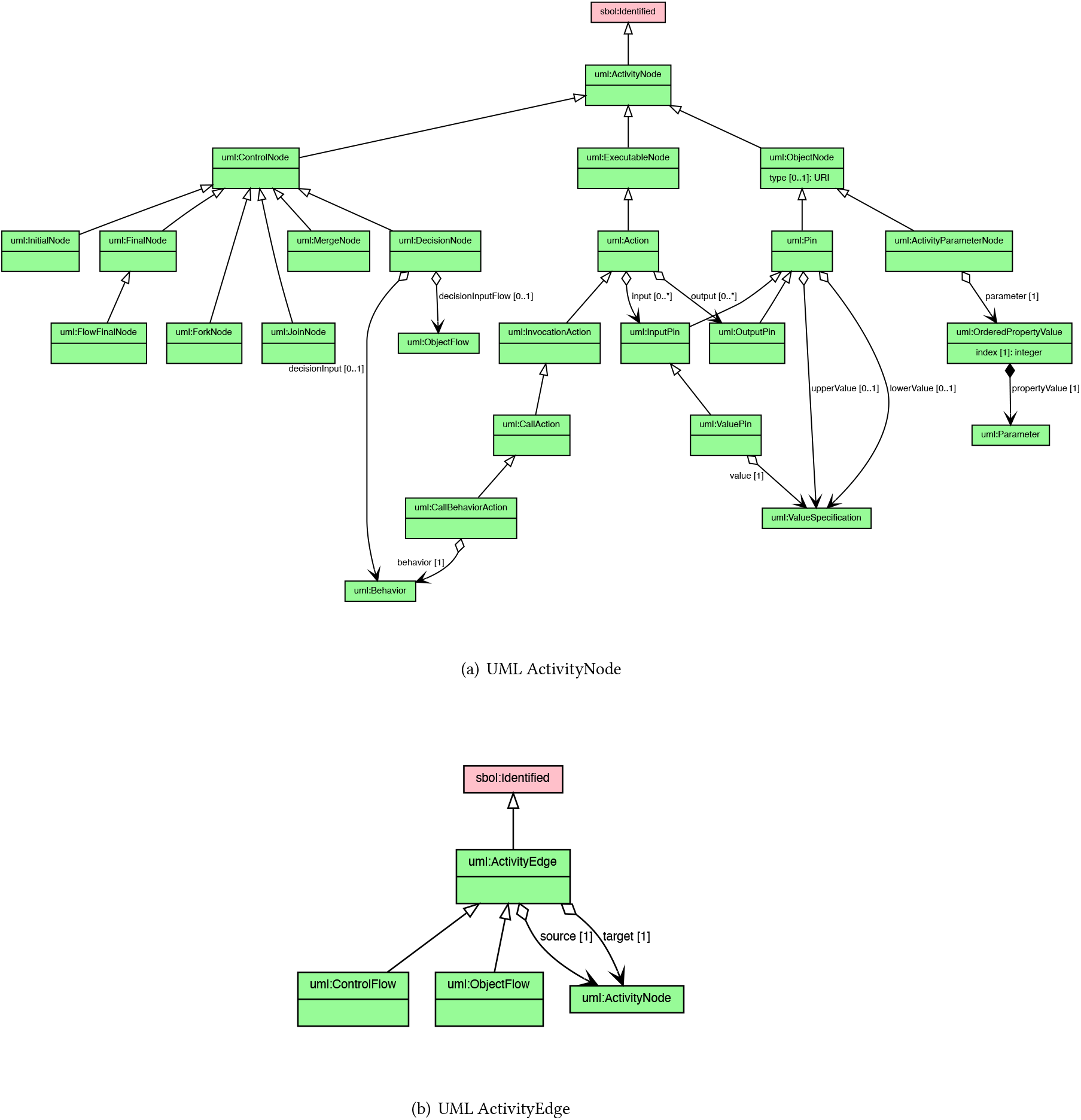
A UML `uml:Activity` is defined as a network of `uml:ActivityNode` objects that define steps, decisions, or values in the activity and `uml:ActivityEdge` objects that connect between them.

The `uml:Activity`Edge objects that connect pairs of `uml:ActivityNode` objects, on the other hand, are much simpler, merely indicating a path by which either a control or object token flows from a source to a target.

The execution semantics for a `uml:Activity` are based on a notion of token flow similar to Petri nets [21]. A Petri net is a graph in which nodes have input edges and output edges. Roughly speaking, a Petri net node is enabled to “fire,” sending a token out all of its output edges, when all of its input edges are filled with tokens; at the same time, those input tokens are removed. When a `uml:Activity` (in this case, the node corresponding to the `paml:Protocol`) begins execution, tokens start in any `uml:InitialNode` or input `uml:ActivityParameterNode`. From there, they flow along the connected `uml:ActivityEdge` for each such source node to the connected target node. As tokens arrive at a `uml:ActivityNode`, it waits until a token has arrived along every edge for which it is a target; once all edges have delivered a token, the `uml:ActivityNode` takes action if needed (i.e., it is a `uml:CallBehaviorAction`), then sends a token along every edge for which it is a source. The exceptions are `uml:DecisionNode`, whose purpose is to choose one of several edges on which to send a token, and `uml:MergeNode`, its complement whose purpose is to accept a token arriving from one of several edges. A `uml:CallBehaviorAction` node, on the other hand, also waits for incoming tokens to the `uml:InputPin` objects that define values for the input parameters of the `uml:Behavior` it will call and sends tokens from its associated `uml:OutputPin` objects. This process continues, with tokens proceeding along nodes and edges until execution comes to an end as tokens are absorbed by `uml:FinalNode` and/or output `uml:ActivityParameterNode` objects (or, if the execution fails, when no further execution is possible because no node has all of its input edges filled).

Token flow implements a universally expressive model of behavior execution. The control flow semantics can support ordered steps, via `uml:ObjectFlow` edges linking pins and `uml:ControlFlow` edges linking steps, as well as steps in parallel or in arbitrary order, via `uml:ForkNode` and a lack of constraining edges. Execution patterns can include loops, by means of `uml:DecisionNode` and circular edge patterns, and recursions, by a `uml:Activity` with a `uml:CallBehaviorAction` that calls itself. Furthermore, since tokens can potentially be communicated between different agents and different types of agent, this model also allows for execution to be distributed, e.g., between several pieces of automation equipment, as a mixture of human and automated execution, or even across a group of collaborating laboratories.

For example, the complete iGEM LUDOX calibration protocol is shown in Figure 5. This `paml:Protcol` consists of 11 `uml:ActivityNode`s (not counting `uml:Pins`) with 15 `uml:ActivityEdges` connecting the nodes into a network. Together, they implement a `uml:Behavior` (in this case, a `paml:Protocol`) with an interface of one input `uml:Parameter`—the wavelength to be measured—and one output `uml:Parameter`, the absorbance measured at the plates.

**Fig. 5.**
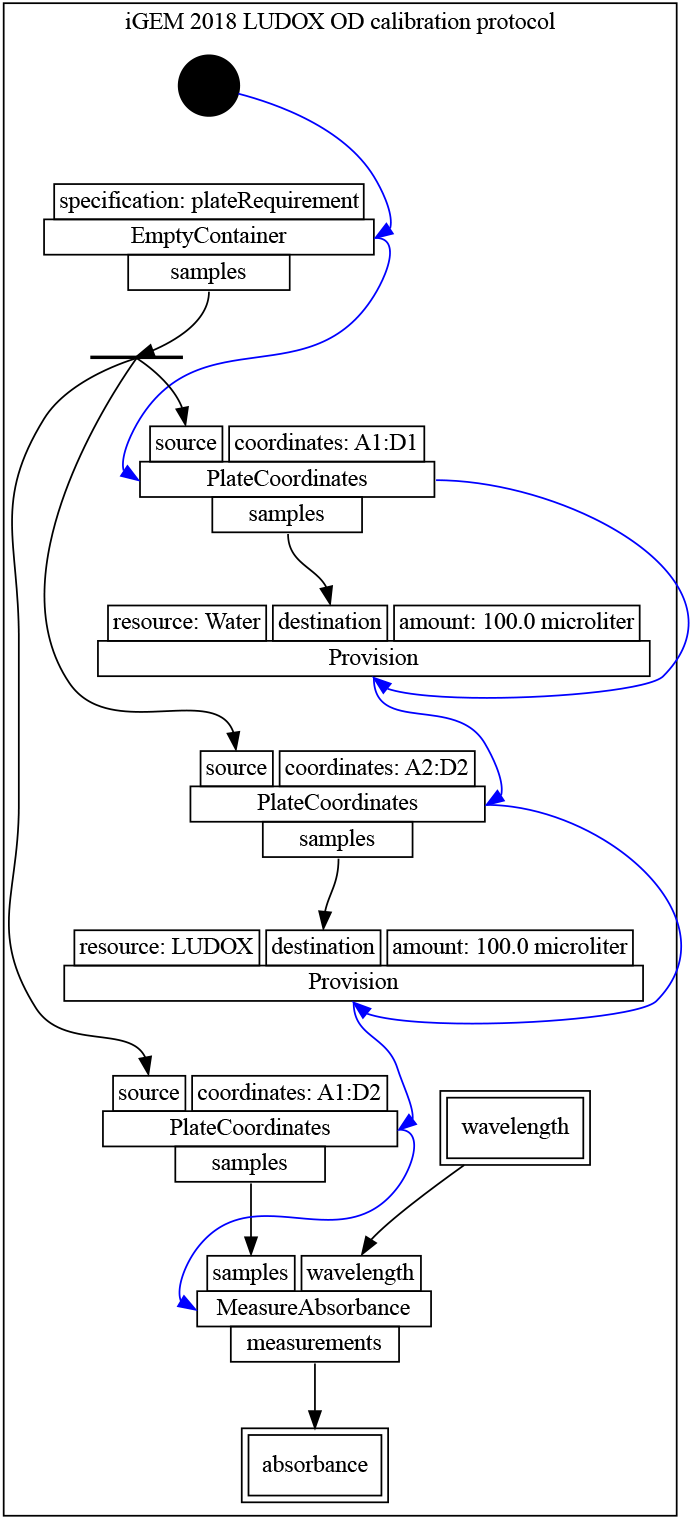
PAML prototocol for the iGEM 2018 LUDOX calibration protocol, automatically rendered by PAML visualizer. The graph includes protocol activities that follow a control flow denoted by blue edges and data flow denoted by black edges. Per UML diagram conventions, the `uml:InitialNode` is shown as a black circle and `uml:ForkNode` is shown as a black bar. The graph also illustrates protocol input (e.g., wavelength) and output (e.g., abosorbance) parameters with double boxes.

In this implementation, two nodes initiate the execution of the protocol: the `uml:InitialNode` and the `uml:ActivityParameterNode` for the wavelength input parameter. To manage the ordering of the steps, `uml:ControlFlow` edges are added, the first of which runs from the `uml:InitialNode` to a `uml:CallBehaviorAction` invoking EmptyContainer from the sample_arrays library. This `paml:Primitive` requests allocation of a `paml:SampleArray` for a 96-well clear flat-bottom plate (specified by the plateRequirement input not expanded in the visualization). Performing `paml:EmptyContainer` in turn enables three `uml:CallBehaviorAction` nodes for PlateCoordinates operations from the same library, with a `uml:ForkNode` implicitly inserted to support the same plate being accessed multiple times.

Note that the Provision operations do not produce outputs, instead acting by modifying state of the `paml:SampleCollection` provided by their destination input. As such, it is necessary to add control edges that ensure that the water is added before the LUDOX, and that both have been put in the plate before absorbance is measured. Finally, once the lab work has been carried out, the final output `uml:ActivityParameterNode` collects and reports the `paml:SampleData` that is output from the call to MeasureAbsorbance.

Since a `paml:Protocol` is itself also a `uml:Behavior`, it can be embedded in other, more complex protocols, including ones with complex hierarchical structures. For example, a `paml:Protocol` might specify a cell culturing protocol that invokes media adjustment and data collection sub-protocols at various time points, or a multi-stage DNA assembly protocol involving multiple rounds of digestion, ligation, transformation, and selection. Furthermore, requesting the execution of a `paml:Protocol` on a particular set of samples or conditions can itself be represented as a simple “wrapper” `paml:Protocol` in which the desired sample and condition values are supplied as `uml:ValuePin` inputs (a `uml:ValuePin` is a `uml:Pin` with a constant value) on a `uml:CallBehaviorAction` invoking the `paml:Protocol` and its results are collected to output via a `uml:ActivityParameterNode` for each expected result. In principle, even an entire experimental campaign could be encoded in such a manner, if desired.

### 3.2 Execution Records

PAML’s representation for recording the trace of an execution is built on the W3C provenance ontology (PROV-O) [18], which provides a mechanism for recording traces. Using this representation will allow PAML to record how data sets are produced as a result of a behavior execution.

PROV-O has its own distinct notion of a `prov:Activity`, in this case a record of an instance of an execution. The basic notion is thus that an execution trace consists of a structure of `prov:Activity` objects, each linked by its `prov:type` property to a corresponding `uml:Behavior` object specifying the nature of the activity. The `prov:Activity` is only a stub class intended for extension, however. Accordingly, in order to provide additional information needed for recording information about protocol executions, PAML extends `prov:Activity` with a `paml:BehaviorExecution` class for recording execution of a UML `uml:Behavior` (either `paml:Primitive` or `paml:Protocol`) and a child `paml:Protocol`Execution class for recording further information about the execution of a `uml:Activity` (i.e., `paml:Protocol`), as shown in Figure 6.

**Fig. 6.**
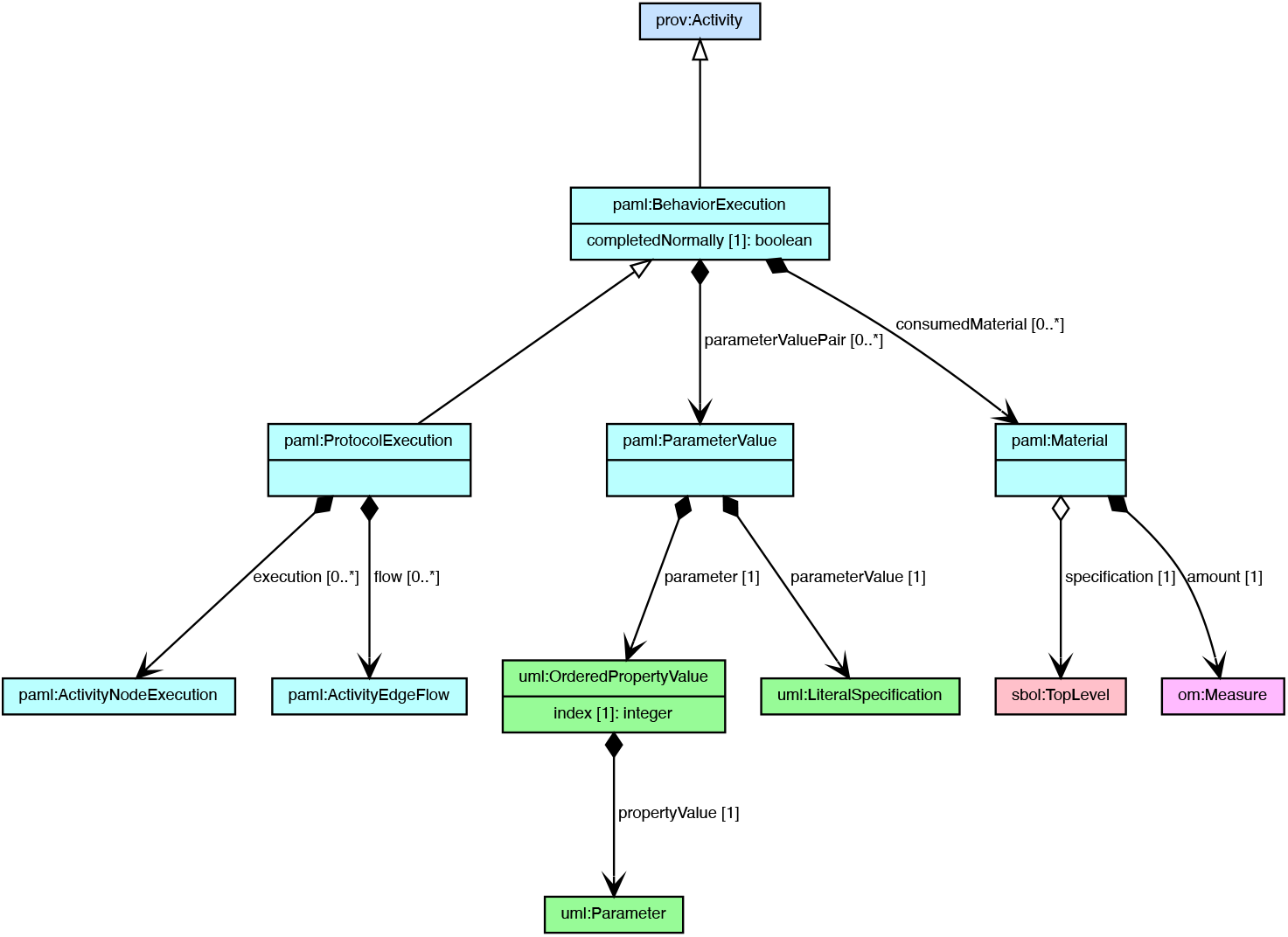
The `paml:Protocol`Execution and `paml:BehaviorExecution` classes are used for recording the execution of `paml:Protocol`s and `paml:Primitives`.

A `paml:BehaviorExecution` is a record of how a `paml:Protocol`, `paml:Primitive`, or other `uml:Behavior` was carried out. Such an execution might be either real or simulated, for example as part of “unrolling” a protocol for certain execution environments as described below in Section 4.3. Properties inherited from PROV-O provide the basic structure of the trace: the `prov:type` property links to the `uml:Behavior`, its `prov:startedAtTime` and `prov:endedAtTime` properties record timing information, and the entity carrying out the execution (e.g., a particular person in the lab, a liquid handling robot, a plate reader) is recorded with a `prov:Association` to a `prov:Agent`. For details on usage of PROV-O within an SBOL RDF context, see Appendix A.1 of [3]. Additional PAML-specific properties record the input and output `paml:Parameter` values for the `paml:Behavior`, the laboratory materials consumed (as pairs of an `sbol:Component` specifying material type with an om:Measure), and whether the execution completed normally or if there was some exception condition.

**Fig. 7.**
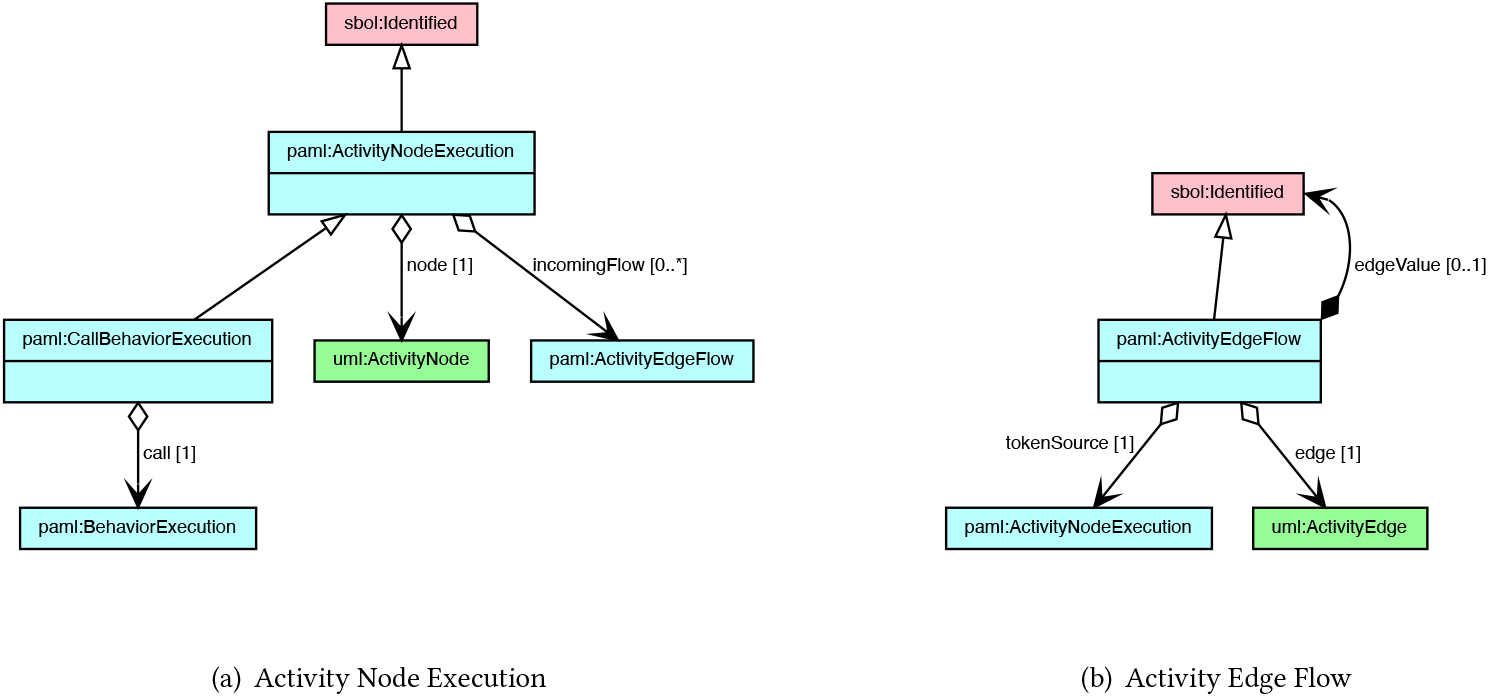
A UML `uml:Activity` is defined as a network of `uml:ActivityNode` objects that define steps, decisions, or values in the activity and `uml:ActivityEdge` objects that connect between them.

A `paml:Protocol`Execution extends this with the addition of records for the nodes and edges defining the `paml:Protocol`’s behavior as a `uml:Activity`. Specifically, the `paml:execution` property is used to record each firing of a `uml:ActivityNode` and the `paml:flow` property is used to record each time a token moves along a `uml:ActivityEdge`, using the `paml:ActivityNodeExecution` and `paml:ActivityEdgeFlow` classes shown in Figure 7. In the case of invocation of a `paml:Primitive` or `paml:Protocol` via a `uml:CallBehaviorAction`, a corresponding `paml:CallBehaviorExecution` subclass of `paml:ActivityNodeExecution` also provides a link down to the `paml:BehaviorExecution` sub-trace that records the execution of the `paml:Primitive` or `paml:Protocol`.

For a simple protocol without any branches or sub-protocols, such as the iGEM LUDOX protocol described above, there will be precisely one execution for each node or edge. For more complex protocols there may be no executions on branches not taken or multiple executions in the case of looping constructs. In either case, however, this parallel construction provides the necessary representation for recording information about the execution of protocols in the lab.

## 4 PROTOTYPE

Following the data model presented above, we have constructed a prototype implementation of PAML in terms of an ontology, derived specification document and Python code library, and prototype software tools for visualization, editing, and execution of PAML protocols.

### 4.1 Ontology-Based Specification

The PAML data representation is implemented first and foremost as an ontology encoded in the Web Ontology Language (OWL). We leverage this machine-readable specification to make the standards development process more efficient through automation. First, we automatically generate graphical visualizations of the data model which illustrate the classes, properties, and links between classes. Second, we automatically generate the data model portions of the PAML specification document as LaTeX. This human-readable document incorporates the class diagrams as well as explanatory descriptions which are sourced from annotations contained in the original ontology file, plus manually generated preamble material introducing the motivation and context of the specification. We also automatically generate an object-oriented Python API which supports authoring and exchange of PAML protocols as any of a number of standard RDF formats. For this purpose, we use a tool called SBOLFactory [4] to dynamically generate Python classes directly from the ontology file.

We use ontology-based specification for several reasons. First, because the standard is still in an early stage of development and is expected to evolve, revisions to the proposed data model can be rapidly generated, released, and tested in practice. Moreover, since the human-readable specification document and the software library are generated from a single source, we avoid introducing errors and discrepancies between the different artifacts. Overall, these factors enable rapid development and responsiveness of PAML developers to emerging needs and use cases from the open protocols community.

The Python API also performs validation on protocols to ensure that representations are complete and consistent. For example, one such validation rule requires that every ProtocolExecution must link to a Protocol. We use the Shapes Constraint Language (SHACL), an RDF-based language that describes graph patterns to which valid data instances must conform [13], to encode these validation rules in a declarative syntax. The pySHACL validator tool [26] can then by used to check conformance of PAML objects in an RDF document according to the encoded rules. By using this combination of OWL and SHACL, we can fully specify the PAML data model using non-ambiguous, machine-readable languages. The specification is thus decoupled from its implementation in any one programming language as well as its formulation in natural language.

### 4.2 Visualization and Editing of Protocols

Graphs are a natural model for visualization and editing of PAML protocols because of their basis in the UML Activity model, in which each protocol involves a set of activities and controls (nodes) that are linked by data and control flows (edges). Figure 5 illustrates the rendering of the iGEM 2018 LUDOX calibration protocol via GraphViz [10]. It includes an initial control node (filled black circle) that is followed by (denoted by a blue control edge) an activity node EmptyContainer. The activity nodes include input pins (e.g., specification) and output pins (e.g., samples). Data flow edges (denoted by black edges) link the activity pins (e.g., EmptyContainer output samples pin links to the source input pin for PlateCoordinates) either directly or through control nodes such as a fork node (denoted by a black bar). Data flow edges also link protocol input (e.g., wavelength) and output (e.g., absorbance) parameters.

While Figure 5 illustrates the protocol as a graph, PAML can be illustrated in a number of formats. The protocol illustrated by Figure 5 may also be described as a list of activities because the activities are totally ordered. PAML protocols that are partially ordered or include decision nodes can be represented by other paradigms such as block-based programs [15] or visual scripts [1, 9]. More broadly, protocols can be viewed as programs that can be expressed in a programming language or pseudocode.

In addition to visualizing protocols, the same paradigms support editing, such as adding, deleting, or configuring activities. The PAML Python API supports protocol editing operations by providing functions that build a protocol. Protocol generation scripts (e.g., as listed in Figure 8) execute a sequence of API functions that construct the PAML for the iGEM LUDOX calibration protocol. For example, in that script lines 5 to 7 define the protocol and add it to an SBOL3 document (representing the protocol as RDF). Line 10 defines an input parameter called wavelength. Line 13 defines the SBOL3 object for double-distilled water, grounding it in a link to a PubChem identifier. Line 18 defines a microplate object that will hold the samples. Lines 21 to 24 identify the wells that will hold water and provision the water into those wells. Lines 29 to 32 identify which wells to measure and then measure the absorbance. Finally, lines 35 to 38 define the protocol output parameter for absorbance and link it to the output of the absorbance measurement activity. Visual editors can use these functions to implement the same functionality as the Python script.

**Fig. 8.**
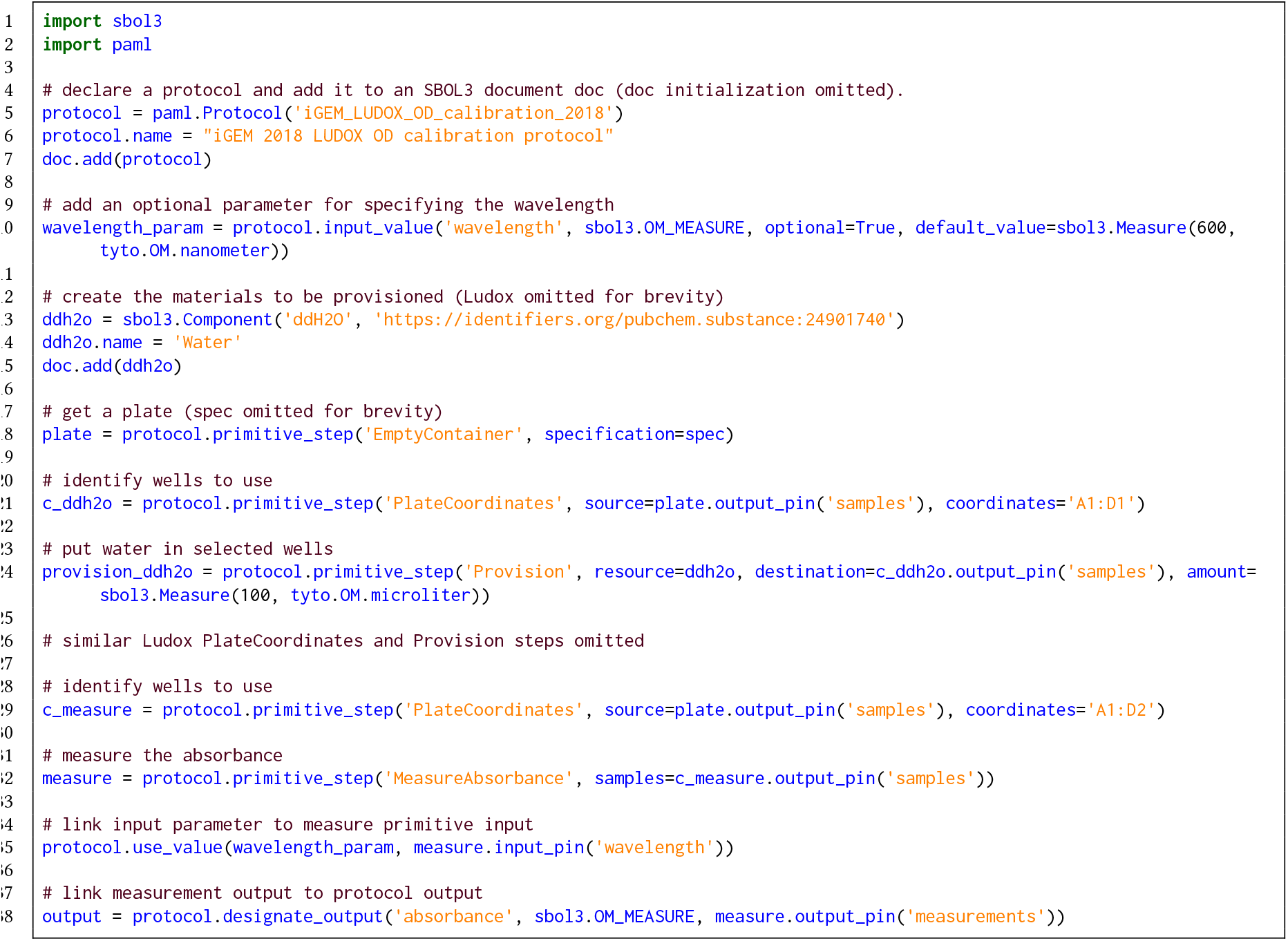
PAML Python script to construct a portion of the iGEM LUDOX calibration protocol. An interactive Jupyter notebook is available at: https://colab.research.google.com/drive/1WPvQ0REjHMEsginxXMj1ewqfFHZqSyM8?usp=sharing

Visualizing protocols, like visualizing source code, helps to specify and understand the protocol. However, like programs, executing a protocol requires interpreting the steps. As the number of objects and control flow nodes increases, it becomes increasingly difficult to interpret the protocol. Furthermore, PAML protocol serialization to different target languages may require compiling away some of the control structure: for example, in Autoprotocol control flow is a strict linear step order with no branching, so a PAML protocol must be linearized with all loops “unrolled” and all sub-protocol invocations expanded inline in order to produce a linearized version of the protocol suitable for mapping into Autoprotocol. Accordingly, in order to support execution either directly or though serialization, we have developed an execution engine for PAML.

### 4.3 Execution in Markdown and Autoprotocol

PAML execution requires interpreting a protocol to determine which activity can be executed next, and then recording the data generated on the output pins. The PAML execution engine uses a token based execution semantics that implements the UML activity model based upon Petri-nets [21]. The execution involves tracking a set of tokens generated by each activity or control node in the protocol. Each activity generates tokens on their output pins and consumes tokens on the input pins. The tokens can either hold data representing objects created by activities, or can denote control. The execution engine non-deterministically selects an enabled protocol node each iteration. It records the execution for each node in an execution trace and notifies any listeners. Executing a protocol offline (e.g., for export to Autoprotocol) involves generating identifiers for activity outputs that will be specified later, during an actual execution of the protocol. Online execution of the protocol provides an opportunity to specify actual outputs as the activities execute. Figure 9 shows an example visualizing an execution trace for the LUDOX protocol.

**Fig. 9.**
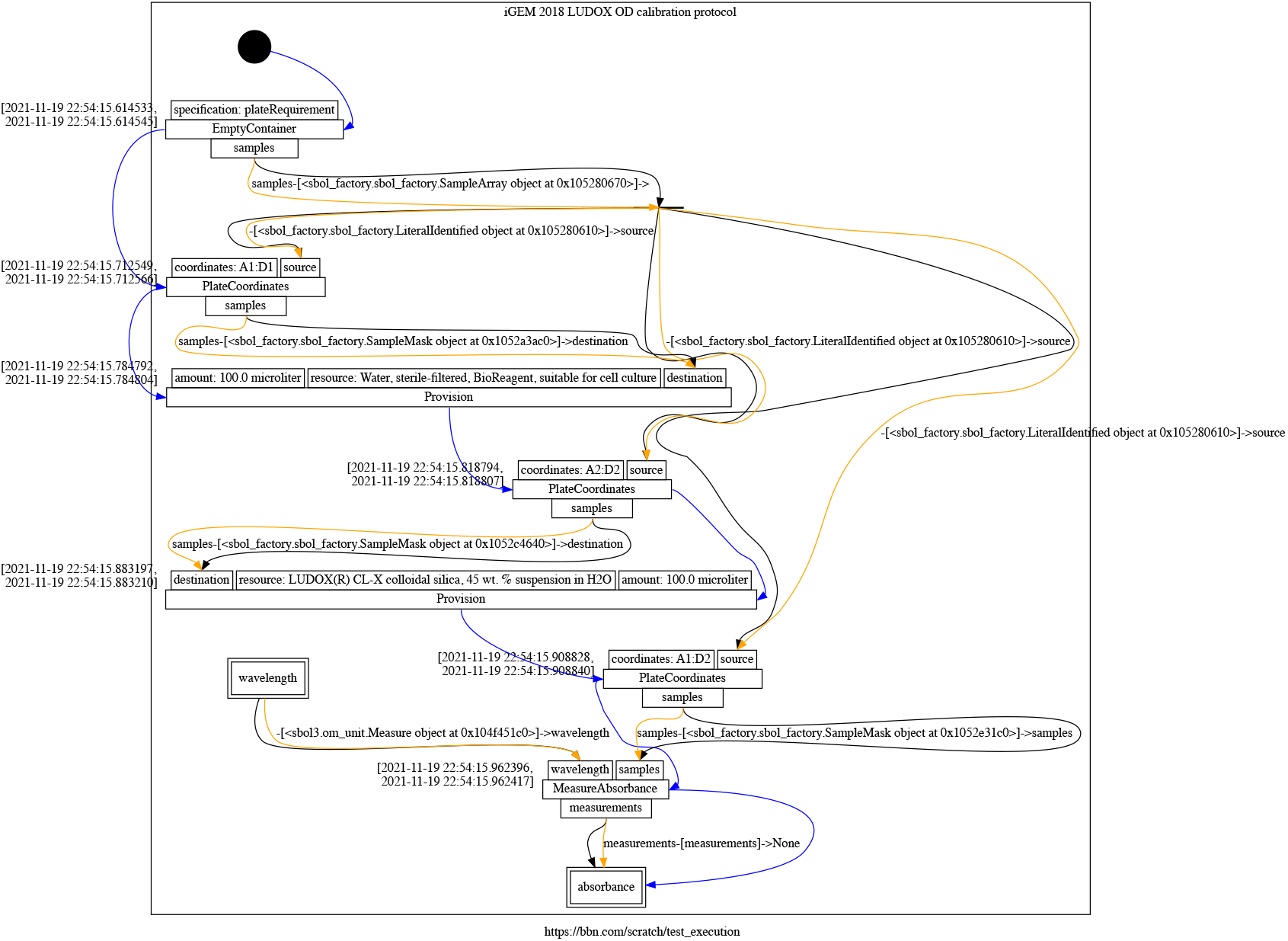
PAML execution trace for the iGEM 2018 LUDOX calibration protocol, layered on the protocol visualization shown in Figure 5. Yellow edges denote data flow with placeholder and computed values for an offline execution of the protocol.

In order to generate serializations of protocols to alternative target formats, the execution engine uses listeners that output either Autoprotocol or Markdown. Both Autoprotocol (a machine language) and Markdown (a list of steps in semi-structured natural language) require a sequence of steps that totally order the protocol activities and omit control flow that is otherwise implicit in the format (e.g., omitting the object fork node for the plate, as illustrated by the black bar in Figure 5). The execution listener pattern is particularly helpful for serializing protocols that include multiple repetitions of sub-protocols as activities. This requires the protocol executor to “unroll” the protocol into a series of distinct executions, in which sub-protocols may appear multiple times as they are repeated.

PAML is converted to Autoprotocol and Markdown with different execution listeners. For example, the iGEM LUDOX calibration protocol is shown converted and rendered into Markdown in Figure 10 and into Autoprotocol in Figure 11. The listeners not only collect the translated sequence of protocol activities, but also help to resolve the objects appearing the protocol. For example, the Autoprotocol listener interprets the EmptyContainer activity to identify an available container (c.f., line 38 in Figure 11) in the Strateos laboratory information management systems (LIMS) that will satisfy the specification made in the protocol. Similarly, the Markdown listener interprets the protocol to construct a human-readable strings that describe each step, as well as embedding links to definitions, for example in this case linking each material to its NCBI PubChem substance definition, which in turn provides supplier information for the required materials. In addition to interpreting the steps, the listeners also format the syntax needed for each target language: the Autoprotocol listener formats protocols as a list of instructions in JSON, and the Markdown listener makes use of Markdown syntax to hyperlink definitions of reagents and containers.

**Fig. 10.**
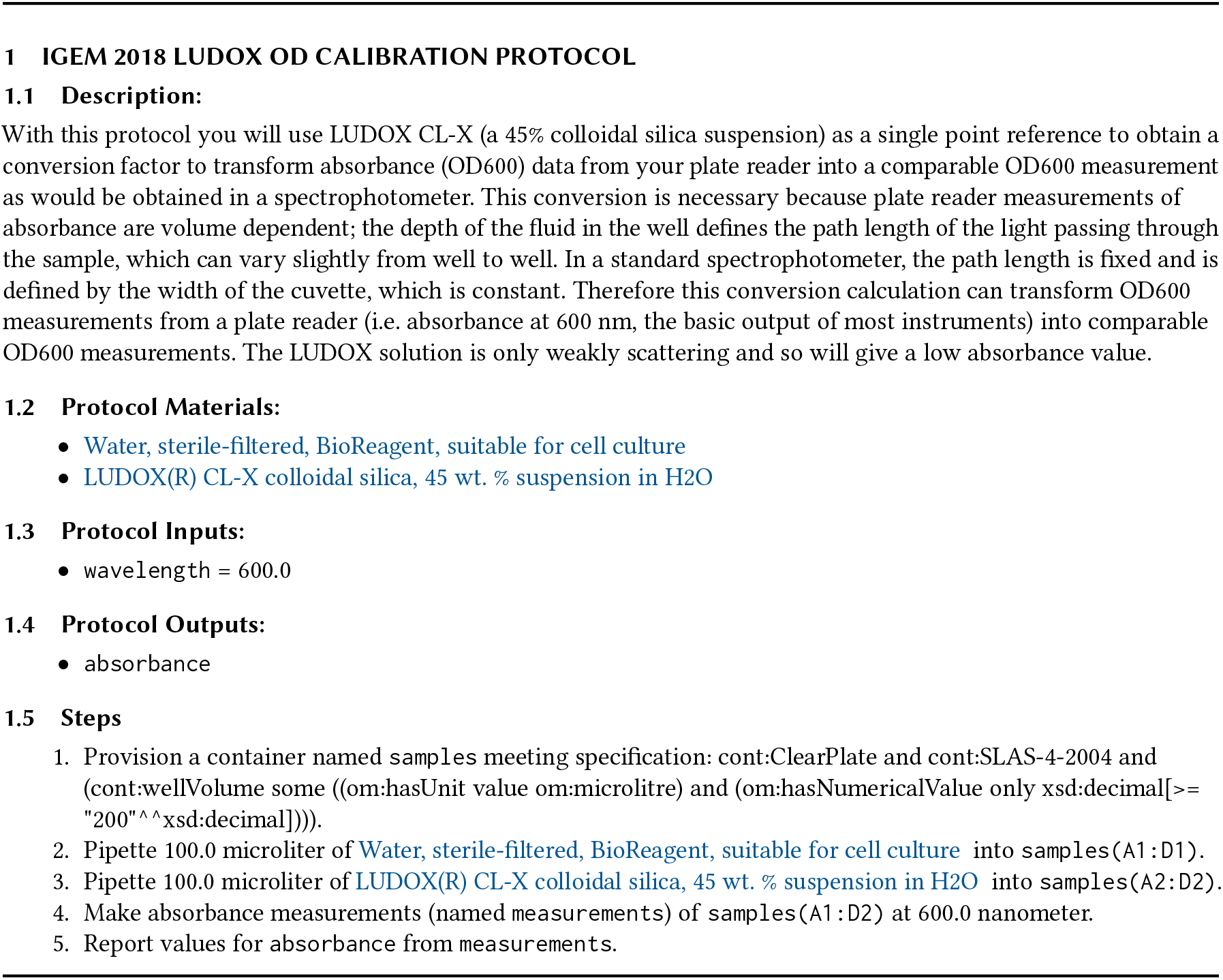
Markdown “paper protocol” generated from PAML for iGEM 2018 LUDOX calibration protocol.

**Fig. 11.**
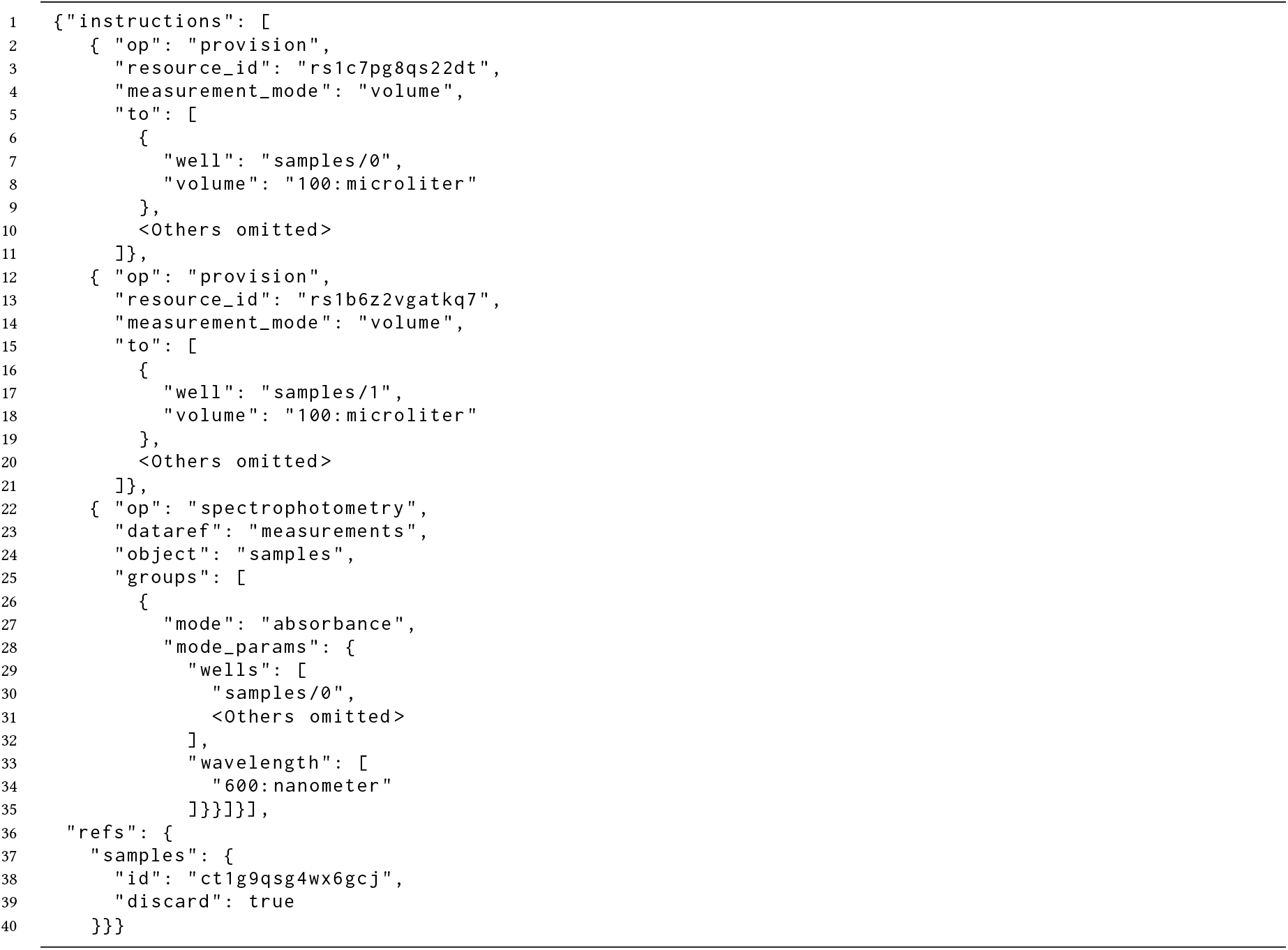
Autoprotocol specification of the iGEM LUDOX calibration protocol generated by the PAML execution engine and Autoprotocol listener.

## 5 FUTURE DIRECTIONS

The development efforts on PAML described above have produced a draft representation that appears to be simultaneously expressive enough and compact enough to satisfy all of the key goals that we have identified for a broadly applicable community standard. Our prototype implementation realizes this representation in the form of an ontology, specification, and Python library, which in turn have been used to implement test protocols and tools for visual editing and for execution, either by hand via export to a “paper protocol” or with laboratory robotics via export to Autoprotocol.

The next critical stage in developing this into an effective community standard for protocols is to refine the representation and expand the set of tools through involvement of interested stakeholders from the broader community. To that end, we organized an open community meeting at the COMBINE 2021 standards meeting in October, 2021, during the course of which participants validated community interest in this initiative, prioritized next steps for PAML, and began organization of an open pre-competitive community for its continued development, which may be found at http://bioprotocols.org/

The key near-term goals for the development of PAML, as currently prioritized by this community, are thus:

- putting PAML to use in ongoing interlaboratory collaborations within the stakeholder community,
- implementation of additional key execution environments, such as the OpenTrons API [20] and protocols.io [27],
- determining representational details for sample arrays and sample data,
- implementing reasoning about the contents of samples, and
- improved user interfaces for protocol design, editing, and inspection.

If this nascent community is able to achieve these goals, particularly in using PAML to reduce the protocol-related challenges faced by existing interlaboratory collaborations, then it will form the basis for further development and utilization and, ultimately, may be able to establish an effective open standard representation for biological protocols, accelerating research and development across a broad range of fields and applications.

## ACKNOWLEDGMENTS

This work was supported by Air Force Research Laboratory (AFRL) and DARPA contracts FA8750-17-C-0184, FA8750-17-C-0231, and HR001117C0095. This document does not contain technology or technical data controlled under either U.S. International Traffic in Arms Regulation or U.S. Export Administration Regulations. Views, opinions, and/or findings expressed are those of the author(s) and should not be interpreted as representing the official views or policies of the Department of Defense or the U.S. Government. Approved for Public Release, Distribution Unlimited.

